## Supplementary Information for "Structural-Functional Brain Network Coupling During Task Performance Reveals Intelligence-Relevant Communication Strategies"

Department of Psychology I

Würzburg D-97070, Germany

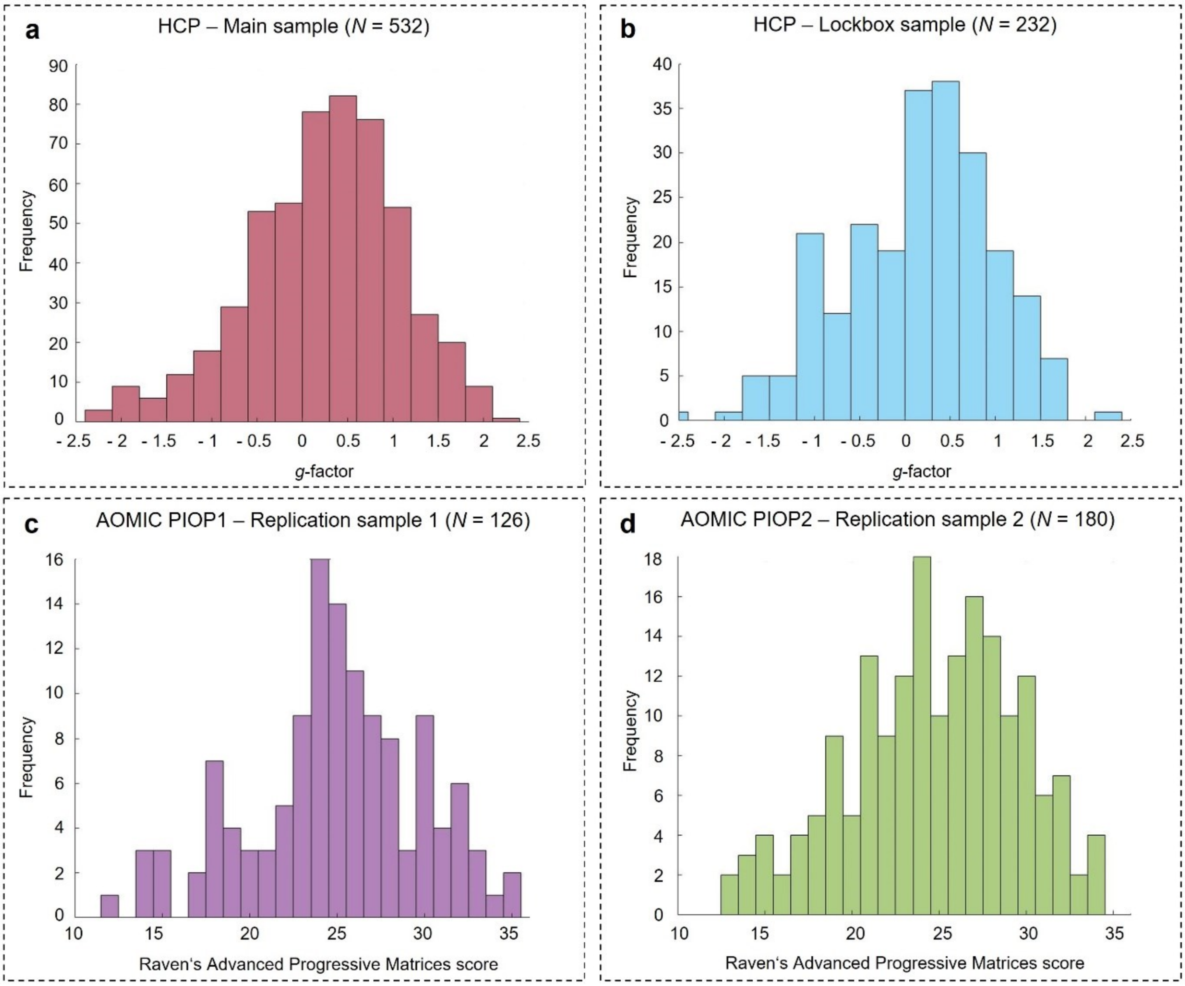


**Supplementary Fig. S1 | Distribution of general intelligence scores in the four samples. a,** In the main sample (HCP; *N* = 532), general intelligence scores operationalized as latent *g*-factor ranged between -2.38-2.40 (mean = .25; standard deviation = .83). **b,** In the lockbox sample (HCP; *N* = 232), general intelligence scores also operationalized as latent *g*-factor ranged from -2.43-2.12 (mean = .15; standard deviation = .83). **c,** In the replication sample AOMIC PIOP1 (*N* = 126) general intelligence was approximated with the sum score of the Raven's Advanced Progressive Matrices Test ranging from 12-35 (mean = 24.90; standard deviation = 4.87). **d,** In the replication sample AOMIC PIOP2 (*N* = 180) general intelligence was also approximated with the sum score of the Raven's Advanced Progressive Matrices Test ranging from 13-34 (mean = 24.68; standard deviation = 4.82). HCP = Human Connectome Project; AOMIC = Amsterdam Open MRI Collection.

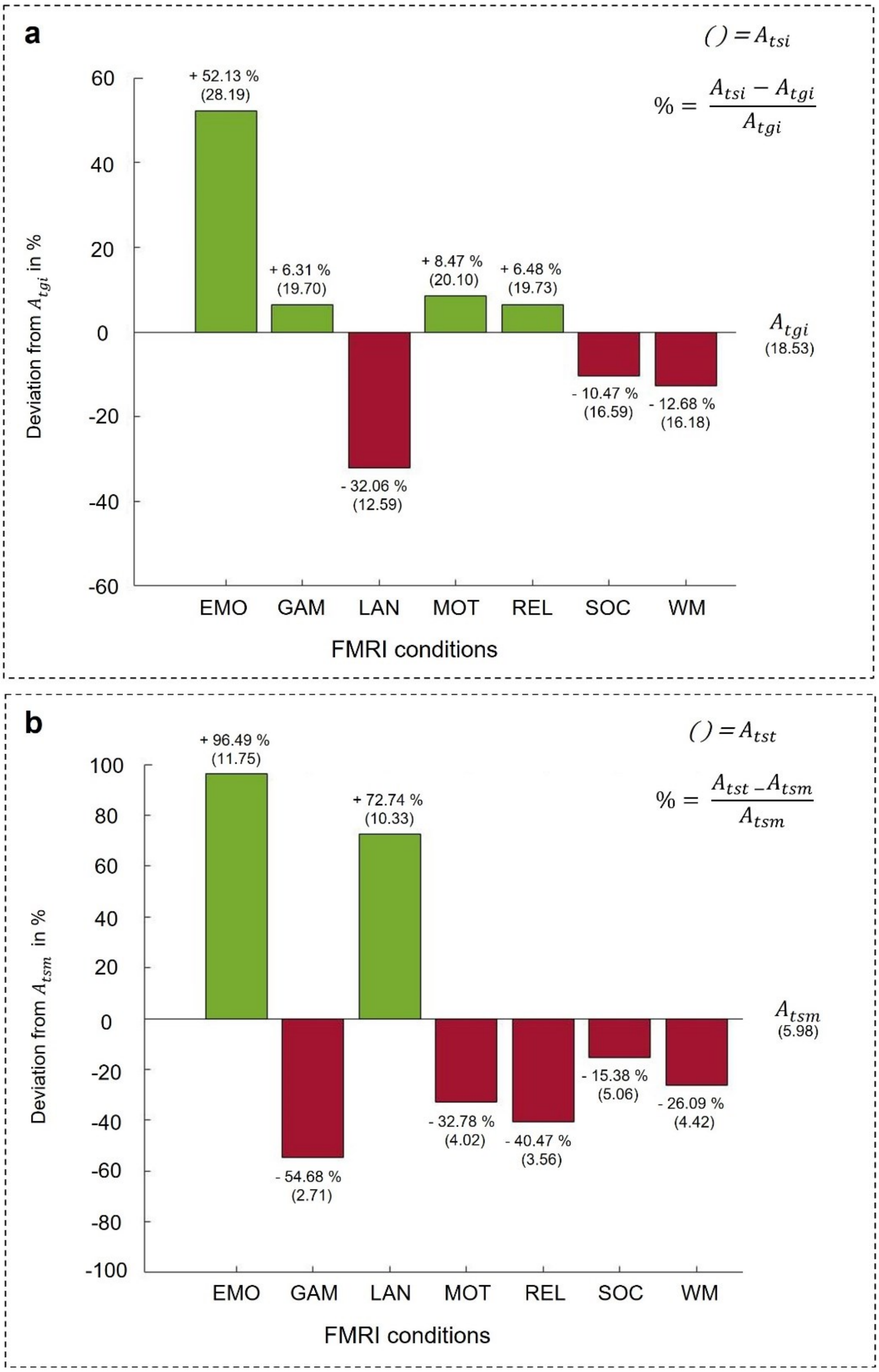


**Supplementary Fig. S2 | Task-specific adaptations from the intrinsic SC-FC coupling pattern and task-general SC-FC coupling patterns**. These were quantified by comparing regional group-average patterns of SC-FC coupling strength. Specifically, the 358-by-1 vectors representing a) the group-average intrinsic SC-FC coupling pattern or b) the group-average task-general SC-FC coupling pattern were subtracted from vectors representing group-average task-specific SC-FC coupling patterns, generating a difference measure quantifying the adaptation from the intrinsic or task-general condition to each specific task fMRI condition. Note that all computations are based on group-general SC-FC coupling patterns. **a,** Absolute task-specific adaptations in region-specific SC-FC coupling from intrinsic condition ($A_{tsi}$, in parentheses) were computed. Here, larger values indicate greater adaptation. These were set in relation to a task-general adaptation from the intrinsic condition ($A_{tgi}$), where larger percentages indicate a greater and lower percentages a smaller adaptation ($\frac{A_{tsi}-A_{tgi}}{A_{tgi}}$). **b,** Absolute task-specific adaptations in region-specific SC-FC coupling from task-general condition ($A_{tst}$, in parentheses) were computed. Again, larger values indicate greater adaptation. They were set in relation to a mean adaptation value computed across all tasks ($A_{tsm}$), where larger percentages indicate a greater and lower percentages a smaller task-specific adaptation ($\frac{A_{tst}-A_{tsm}}{A_{tsm}}$). EMO = emotion processing task; GAM = gambling task; LAN = language task; MOT = motor task; REL = relational processing task; SOC = social cognition task; WM = working memory task.

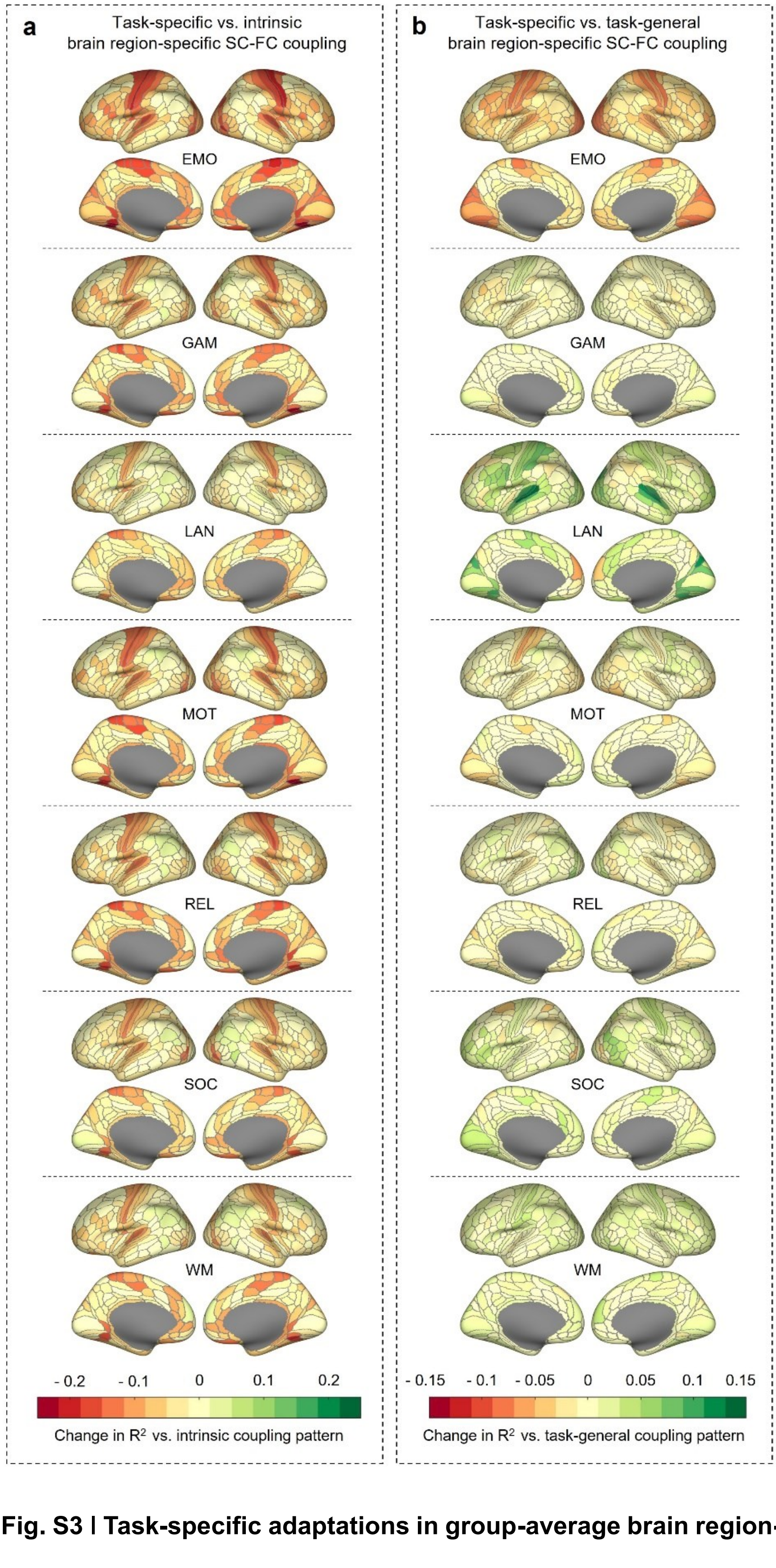


**Supplementary Fig. S3 | Task-specific adaptations in group-average brain region-specific SC-FC coupling patterns. a,** Change in region-specific SC-FC coupling ($R^2$) when comparing group-average task-specific regional SC-FC coupling patterns to the group-average intrinsic SC-FC coupling pattern approximated during resting state. Positive values indicate higher task-specific coupling compared to intrinsic coupling while negative values indicate lower task-specific coupling compared to intrinsic coupling. **b,** Change in region-specific SC-FC coupling ($R^2$) when comparing group-average task-specific regional SC-FC coupling patterns to a group-average task-general SC-FC coupling pattern (average of region-specific SC-FC coupling values across all remaining tasks). Positive values indicate higher task-specific coupling compared to task-general coupling while negative values indicate lower task-specific coupling compared to task-general coupling. RES = resting state; EMO = emotion processing task; GAM = gambling task; LAN = language task; MOT = motor task; REL = relational processing task; SOC = social cognition task; WM = working memory task.

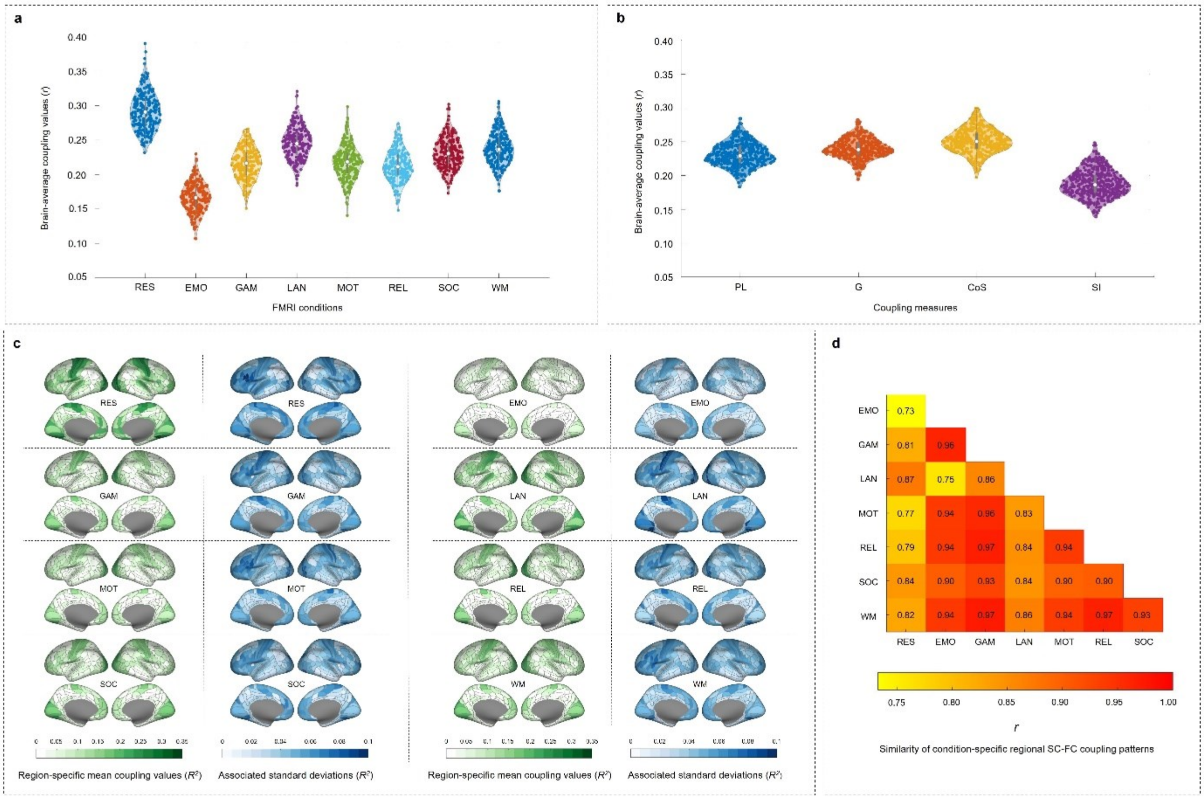


**Supplementary Fig. S4 | Structural-functional brain network coupling varies between measures, fMRI conditions, and participants also in the lockbox sample (HCP232; *N* = 232).** **a,** Distribution of individual condition-specific brain-average SC-FC coupling values. **b,** Distribution of individual measure-specific brain-average SC-FC coupling values. **c,** Condition-specific group-average pattern of SC-FC coupling strength (in green). Blue maps indicate the variance in SC-FC coupling strength (as standard deviation) across participants. **d,** Heatmap depicting the similarity of group-average patterns of brain region-specific SC-FC coupling strength between eight fMRI conditions: Pearson correlation coefficients between condition-specific vectors representing the group-average pattern of SC-FC coupling strength. Note that significant group differences in **a** and **b** are reported in Supplementary Table S6 and Supplementary Table S9, respectively. RES = resting state; EMO = emotion processing task; GAM = gambling task; LAN = language task; MOT = motor task; REL = relational processing task; SOC = social cognition task; WM = working memory task; PL = path length; G = communicability; CoS = cosine similarity; SI = search information.

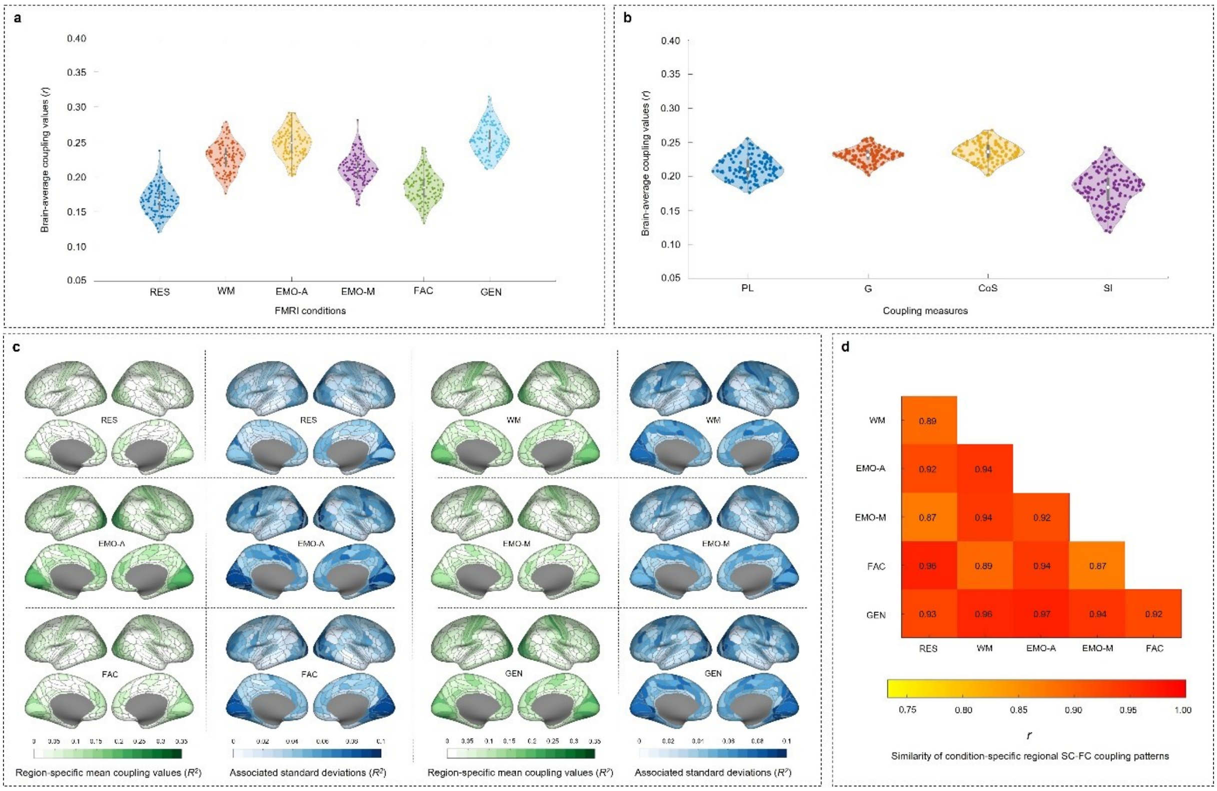


**Supplementary Fig. S5 I Structural-functional brain network coupling varies between measures, fMRI conditions, and participants also in the replication sample (AOMIC PIOP1; *N* = 126). a,** Distribution of individual condition-specific brain-average SC-FC coupling values. **b,** Distribution of individual measure-specific brain-average SC-FC coupling values. **c,** Condition-specific group-average pattern of SC-FC coupling strength (in green). Blue maps indicate the variance in SC-FC coupling strength (as standard deviation) across participants. **d,** Heatmap depicting the similarity of group-average patterns of brain region-specific SC-FC coupling strength between six fMRI conditions: Pearson correlation coefficients between condition-specific vectors representing the group-average pattern of SC-FC coupling strength. Please note that significant group differences in **a** and **b** are reported in Supplementary Table S7 and S10, respectively. RES = resting state; WM = working memory task; EMO-A = emotion anticipation task; EMO-M = emotion matching task; FAC = face perception task; GEN = gender stroop task; PL = path length; G = communicability; CoS = cosine similarity; SI = search information.

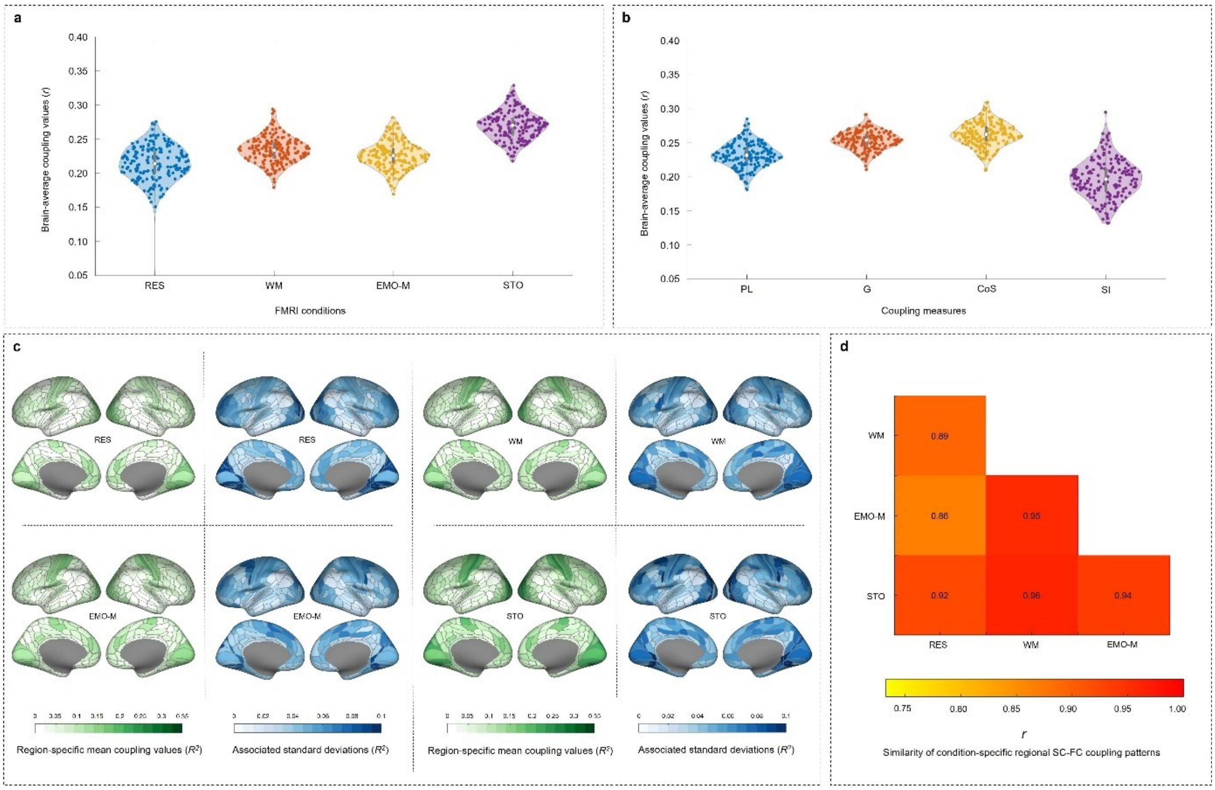


**Supplementary Fig. S6 | Structural-functional brain network coupling varies between measures, fMRI conditions, and participants also in the replication sample (AOMIC PIOP2; *N* = 180). a,** Distribution of individual condition-specific brain-average SC-FC coupling values. **b,** Distribution of individual measure-specific brain-average SC-FC coupling values. **c,** Condition-specific group-average pattern of SC-FC coupling strength (in green). Blue maps indicate the variance in SC-FC coupling strength (as standard deviation) across participants. **d,** Heatmap depicting the similarity of group-average patterns of brain region-specific SC-FC coupling strength between four fMRI conditions: Pearson correlation coefficients between condition-specific vectors representing the group-average pattern of SC-FC coupling strength. Please note that significant group differences in **a** and **b** are reported in Supplementary Table S8 and Supplementary Table S11, respectively. RES = resting state; WM = working memory task; EMO-M = emotion matching task; STO = stop signal task; PL = path length; G = communicability; CoS = cosine similarity; SI = search information.

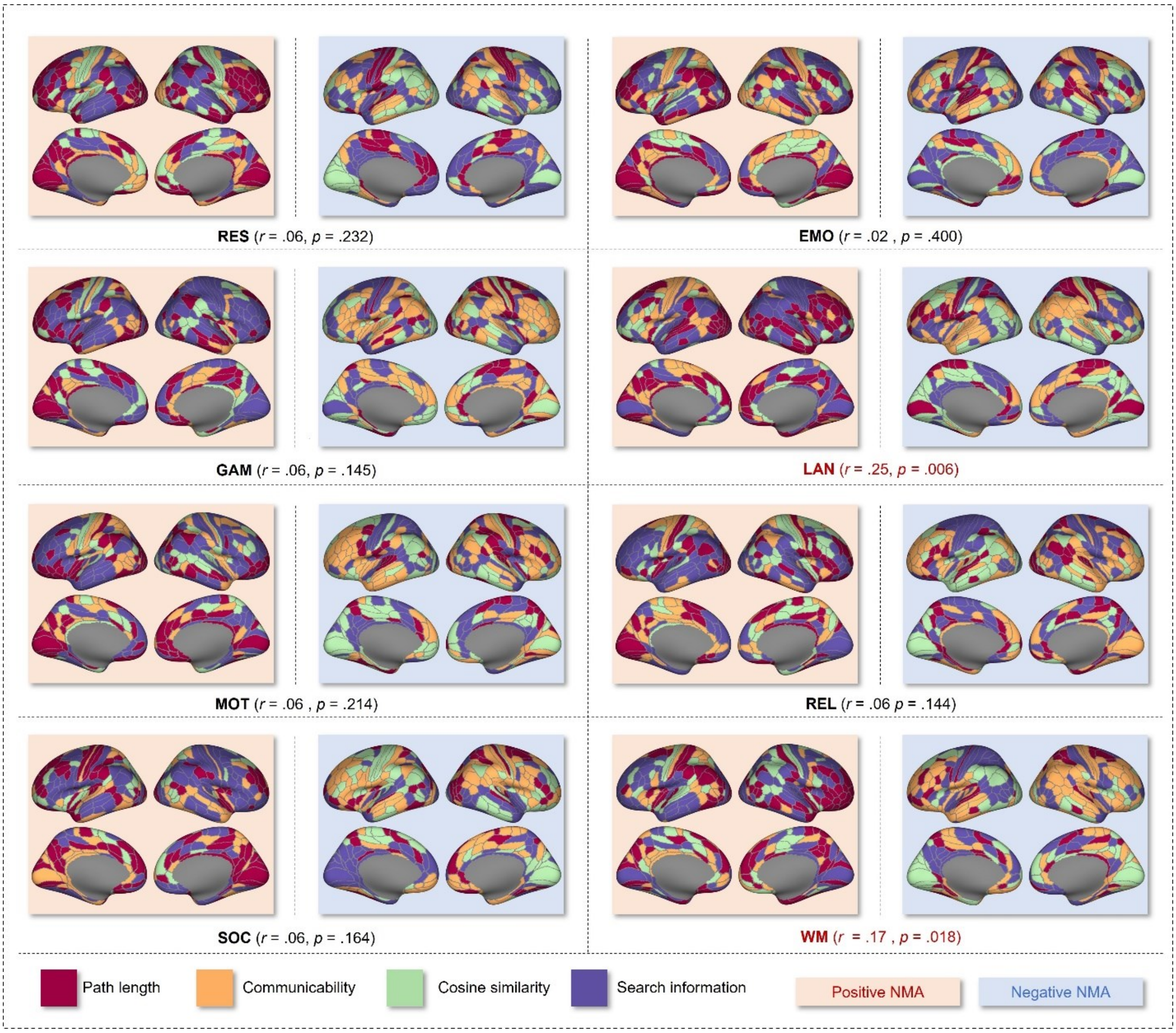


**Supplementary Fig. S7 l Brain region-specific communication strategies in the lockbox sample (HCP232; *N* = 232).** Assignments of coupling measures to brain regions (node-measure assignments; NMAs) were defined by the largest positive (red background) and negative (blue background) magnitude associations between coupling measures and general intelligence across participants. Assignments were created for each fMRI condition separately (Basic NMA Model). The associated prediction performance (correlation between predicted and observed intelligence scores) and significance are reported in parentheses. Note, that the here visualized NMAs are based on data from the whole lockbox sample, whereas the NMAs actually applied in the prediction models were cross-validated, i.e., only derived from data of the training samples. RES = resting state; EMO = emotion processing task; GAM = gambling task; LAN = language task; MOT = motor task; REL = relational processing task; SOC = social cognition task; WM = working memory task.

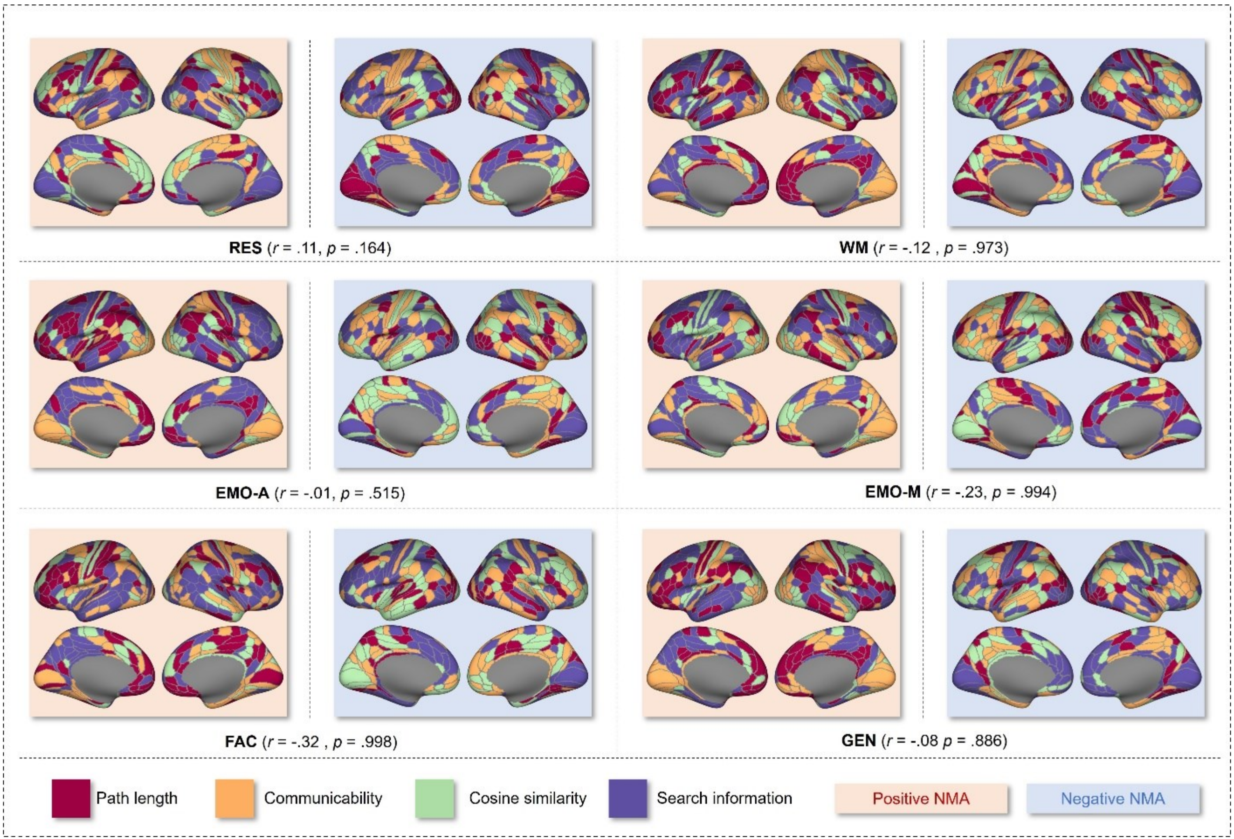


**Supplementary Fig. S8 I Brain region-specific communication strategies in the replication sample (AOMIC PIOP1; *N* = 126).** Assignments of coupling measures to brain regions (NMAs) were defined by the largest positive (red background) and negative (blue background) magnitude associations between coupling measures and general intelligence across participants. Assignments were created for each fMRI condition separately (Basic NMA Model). The associated prediction performance (correlation between predicted and observed intelligence scores) and significance are reported in parentheses. Note, that the here visualized NMAs are based on data from the whole AOMIC PIOP1 sample, whereas the NMAs actually applied in the prediction models were cross-validated, i.e., only derived from data of the training samples. RES = resting state; WM = working memory task; EMO-A = emotion anticipation task; EMO-M = emotion matching task; FAC = face perception task; GEN = gender stroop task; PL = path length; G = communicability; CoS = cosine similarity; SI = search information.

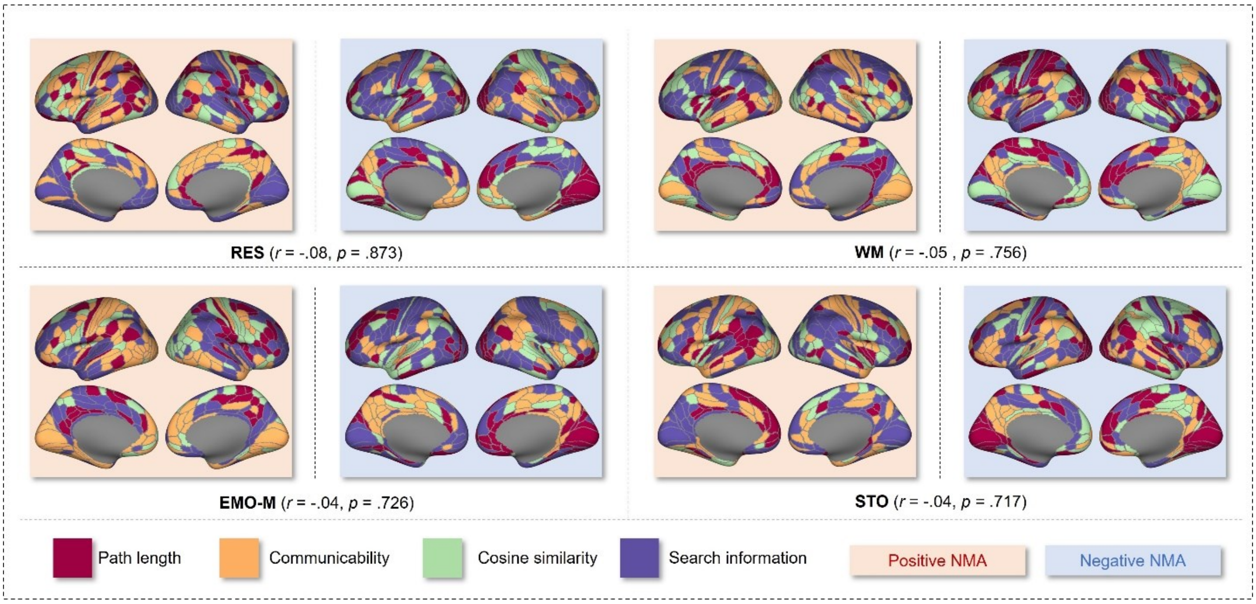


**Supplementary Fig. S9 | Brain region-specific communication strategies in the replication sample (AOMIC PIOP2; $N$ = 180).** Assignments of coupling measures to brain regions (NMAs) were defined by the largest positive (red background) and negative (blue background) magnitude associations between coupling measures and general intelligence across participants. Assignments were created for each fMRI condition separately (Basic NMA Model). The associated prediction performance (correlation between predicted and observed intelligence scores) and significance are reported in parentheses. Note, that the here visualized NMAs are based on data from the whole AOMIC PIOP2 sample, whereas the NMAs actually applied in the prediction models were cross-validated, i.e., only derived from data of the training samples. RES = resting state; WM = working memory Task; EMO-M = emotion matching task; STO = stop signal task.

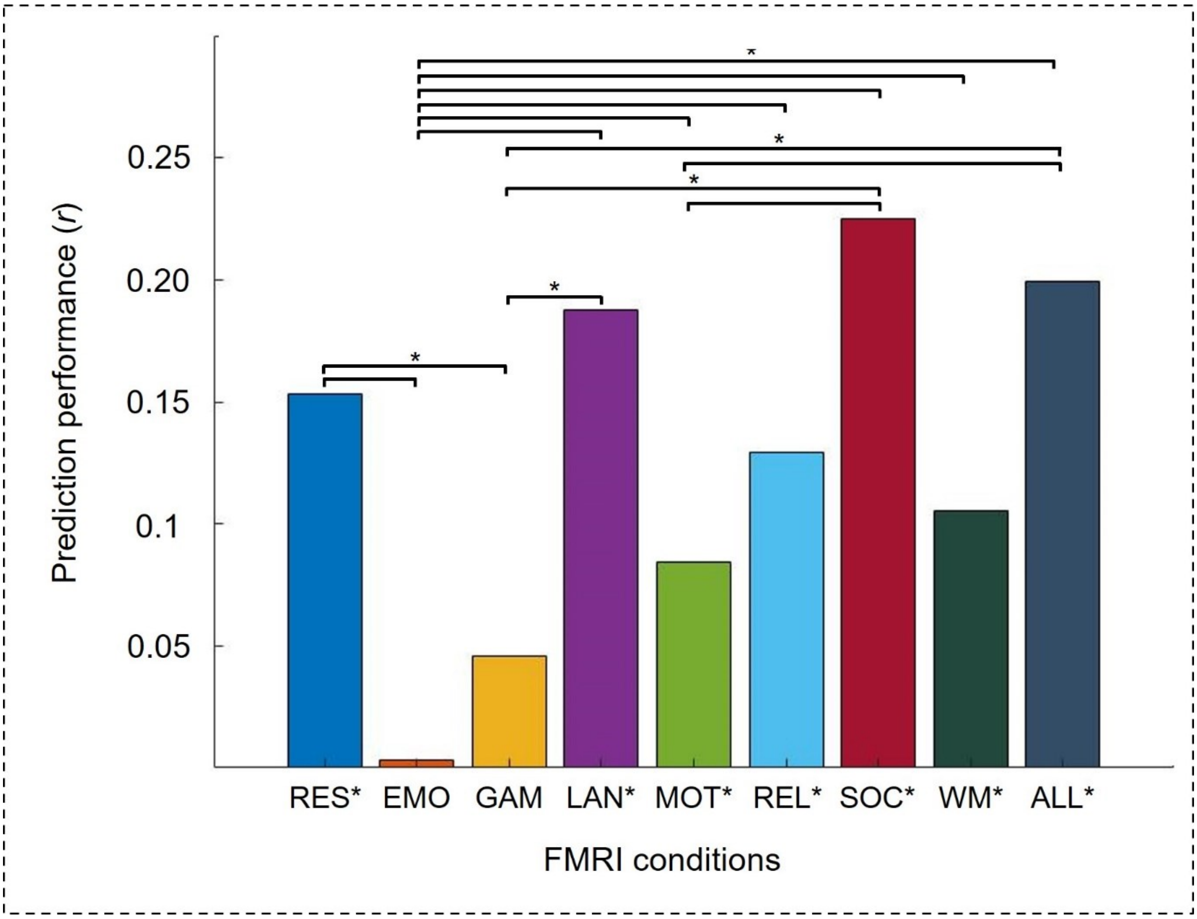


**Supplementary Fig. S10 | Performance to predict intelligence from SC-FC coupling during different fMRI conditions (Basic NMA Model) and a combination of all task fMRI conditions (Expanded NMA Model) in the cross-sample model generalization test in the lockbox sample (HCP532 → HCP232).** The significance of a prediction performance was assessed with a non-parametric permutation test. Significant differences in model performance were assessed with model difference tests (see Methods). *P*-values indicating significant associations are marked with an asterisk a) below bar graphs for single prediction performance and b) above bar graphs for differences in prediction performance (* = $p < .05$). RES = resting state; EMO = emotion processing task; GAM = gambling task; LAN = language task; MOT = motor task; REL = relational processing task; SOC = social cognition task; WM = working memory task; ALL = all task fMRI conditions.

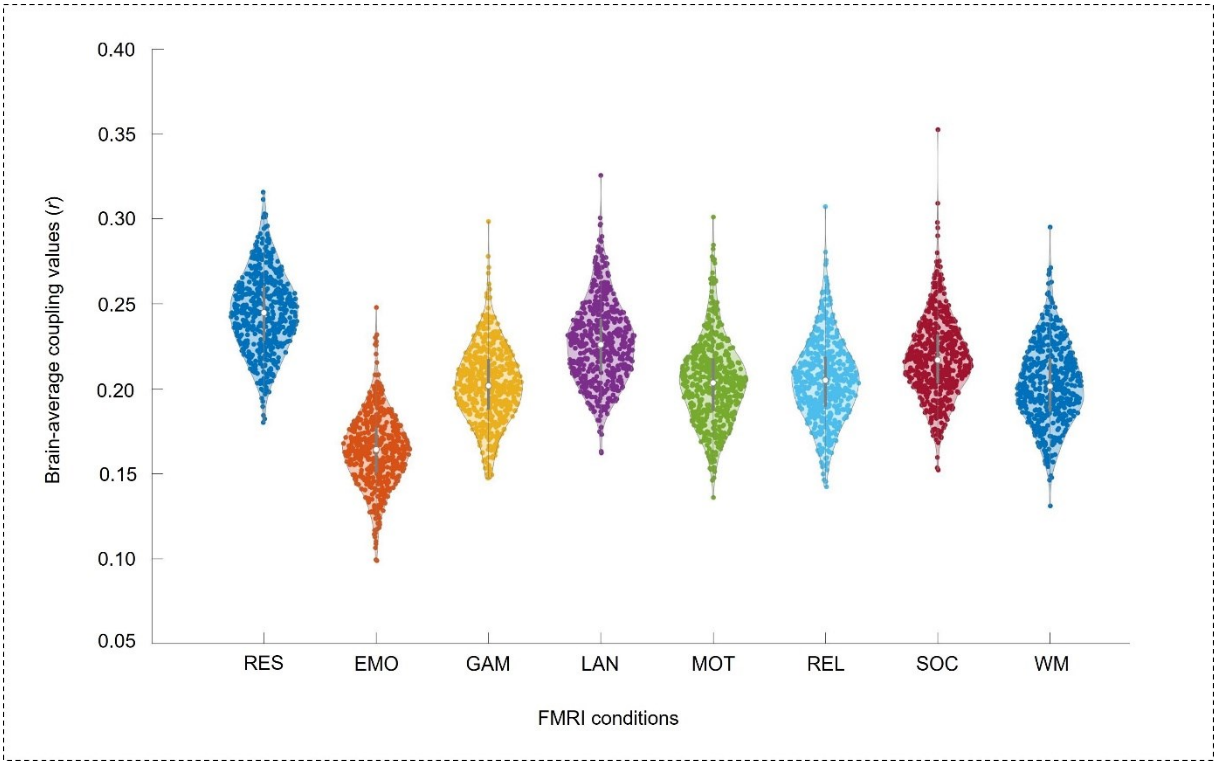


**Supplementary Fig. S11 I Control analysis equal frame length.** Distribution of individual measure-specific brain-average SC-FC coupling values when all scan lengths are trimmed to a frame length of 176 (matching the shortest task: emotion processing task) prior to computing the region-specific SC-FC coupling values. RES = resting state; EMO = emotion processing task; GAM = gambling task; LAN = language task; MOT = motor task; REL = relational processing task; SOC = social cognition task; WM = working memory task; ALL = all task fMRI conditions.

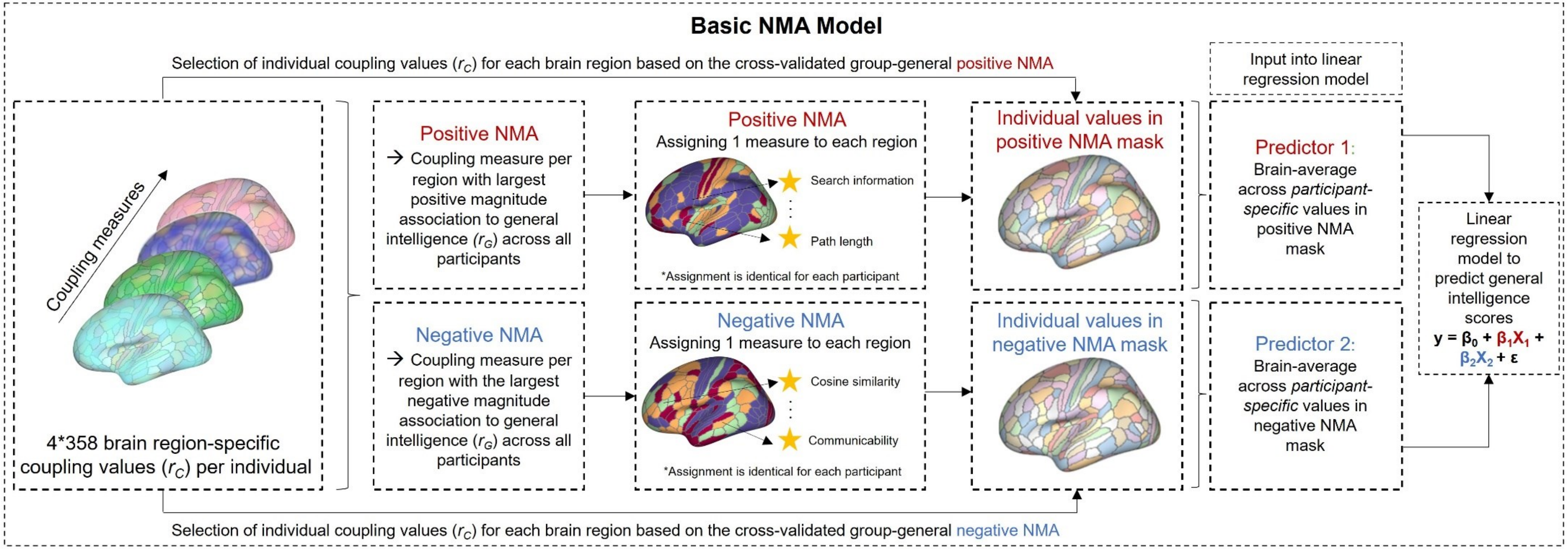


**Supplementary Fig. S12 | Visualization of the Basic NMA prediction model used to predict individual general intelligence from region-specific SC-FC coupling.** Two global input features were generated by first determining a group-general positive and negative node-measure assignment mask (NMA) designating the measure with the largest positive magnitude association (positive NMA) and the largest negative magnitude association (negative NMA) to a given brain region. These group-general masks were used to extract participant-specific regional coupling values ($r_C$) and ultimately, brain-averages across the participant-specific values in the positive NMA mask (predictor 1) and in the negative NMA mask (predictor 2) served as the two predictors for the 5-fold cross-validated multiple linear regression model. Importantly, NMAs built in the training samples were used to extract region-specific coupling values in the respective test samples to avoid data leakage between cross-validation folds. Prediction performance was assessed by correlating predicted and observed general intelligence scores and significance was determined with non-parametric permutation tests. NMA = node-measure assignment.

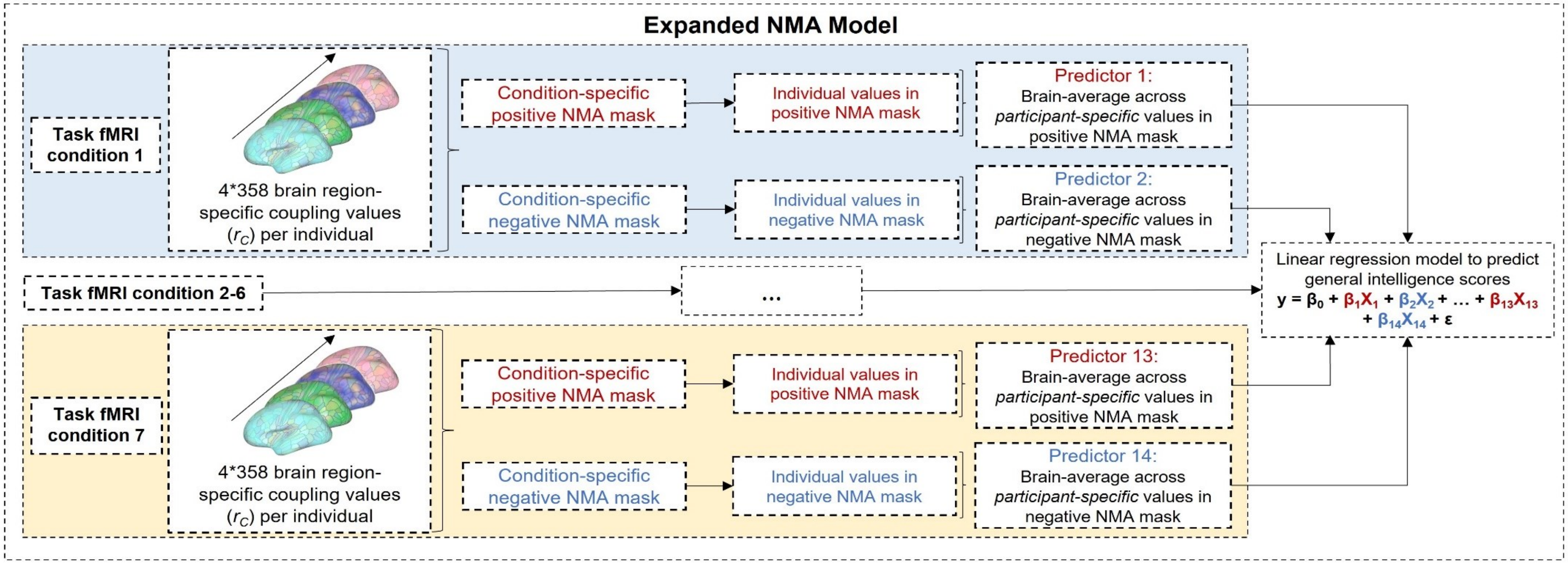


**Supplementary Fig. S13 I Visualization of the Expanded NMA prediction model used to predict individual general intelligence from region-specific SC-FC coupling.** For each task fMRI condition separately, group-general NMA masks were created based on the largest positive magnitude association (Positive NMA) and the largest negative magnitude association (Negative NMA) between region-specific coupling measures and general intelligence. These group-general masks were used to extract participant-specific regional coupling values ($r_C$) for each respective task fMRI condition. Ultimately, brain-averages across the participant-specific values in the positive NMA mask (predictor 1) and in the negative NMA mask (predictor 2) per task fMRI condition were used as model inputs, yielding a total of 14 predictors for the 5-fold cross-validated multiple linear regression model. To guarantee thorough cross-validation and prevent data leakage between cross-validation folds, NMAs built in the training samples were used to extract region-specific SC-FC coupling values in the respective test samples.

Prediction performance was assessed by correlating predicted and observed general intelligence scores and significance was determined with non-parametric permutation tests. NMA = node-measure assignment.

**Supplementary Table S1**

Cognitive tests and measures used to calculate a latent *g*-factor as estimate of general intelligence.

| Test | Instrument | Measure used as input for *g*-factor calculation |
|---|---|---|
| **1** | Episodic memory (Picture sequence memory) | PicSeq_Unadj |
| **2** | Executive function/cognitive flexibility (Dimensional change card sort) | CardSort_Unadj |
| **3** | Executive function/inhibition (Flanker task) | Flanker_Unadj |
| **4** | Fluid intelligence (Penn Progressive Matrices) | PMAT24_A_CR |
| **5** | Language/reading decoding (Oral reading recognition) | ReadEng_Unadj |
| **6** | Language/vocabulary comprehension (Picture vocabulary) | PicVocab_Unadj |
| **7** | Processing speed (Pattern completion processing speed) | ProcSpeed_Unadj |
| **8** | Self-regulation/impulsivity (Delay discounting) | DDisc_AUC_200 + DDisc_AUC_40K |
| **9** | Spatial orientation (Variable Short Penn Line Orientation Test) | VSPLOT_TC |
| **10** | Sustained attention (Short Penn Continuous Performance Test) | $\frac{SCPT\_TP + SCPT\_TN}{(SCPT\_TP + SCPT\_TN + SCPT\_FP + SCPT\_FN)SCPT\_TPRT}$ |
| **11** | Verbal episodic memory (Penn Word Memory Test) | IWRD_TOT |
| **12** | Working memory (List sorting) | ListSort_Unadj |

*Note:* Bi-factor analysis was performed to estimate a latent factor of general intelligence[1,2]. Analyses were conducted in a larger sample (*N* = 1186)[3] utilizing data from 12 cognitive measures[4].

**Supplementary Table S2**

Differences in brain-average SC-FC coupling between fMRI conditions.

| FMRI condition 1 | FMRI condition 2 | *p*-value |
|---|---|---|
| RES | EMO | < .001* |
| RES | GAM | < .001* |
| RES | LAN | < .001* |
| RES | MOT | < .001* |
| RES | REL | < .001* |
| RES | SOC | < .001* |
| RES | WM | < .001* |
| EMO | GAM | < .001* |
| EMO | LAN | < .001* |
| EMO | MOT | < .001* |
| EMO | REL | < .001* |
| EMO | SOC | < .001* |
| EMO | WM | < .001* |
| GAM | LAN | < .001* |
| GAM | MOT | .620 |
| GAM | REL | 1.00 |
| GAM | SOC | < .001* |
| GAM | WM | < .001* |
| LAN | MOT | < .001* |
| LAN | REL | < .001* |
| LAN | SOC | < .001* |
| LAN | WM | < .001* |
| MOT | REL | .763 |
| MOT | SOC | < .001* |
| MOT | WM | < .001* |
| REL | SOC | < .001* |
| REL | WM | < .001* |
| SOC | WM | < .001* |

*Note*: Brain-average SC-FC coupling strength significantly differed between fMRI conditions (ANOVA: $F(7,4248) = [1170.4]$, $p < .001$). Significant group differences are marked with an asterisk (Tukey's HSD procedure: * = $p < .05$; uncontrolled for multiple comparisons). RES = resting state; EMO = emotion processing task; GAM = gambling task; LAN = language task; MOT = motor task; REL = relational processing task; SOC = social cognition task; WM = working memory task.

**Supplementary Table S3**

Differences in brain-average SC-FC coupling between coupling measures.

| Measure 1 | Measure 2 | *p*-value |
|---|---|---|
| PL | G | < .001* |
| PL | CoS | < .001* |
| PL | SI | < .001* |
| G | CoS | < .001* |
| G | SI | < .001* |
| CoS | SI | < .001* |

*Note*: Brain-average SC-FC coupling strength significantly differed between coupling measures (ANOVA: F(3,2124) = [1306.2], $p$ < .001). Significant group differences are marked with an asterisk (Tukey's HSD procedure: * = $p$ < .05; uncontrolled for multiple comparisons). PL = path length; G = communicability; CoS = cosine similarity; SI = search information.

**Supplementary Table S4**

Relationship between general intelligence and brain-average SC-FC coupling during all fMRI conditions.

| **SC-FC coupling measure** | **FMRI condition** *r* (*p*) | | | | | | | | | |
|---|---|---|---|---|---|---|---|---|---|---|
| | RES | EMO | GAM | LAN | MOT | REL | SOC | WM | TG | CG |
| **PL** | .05 (.292) | .11 (.011)* | .02 (.617) | -.03 (.442) | -.01 (.886) | .11 (.016) | .09 (.036) | .08 (.071) | .06 (.186) | .06 (.181) |
| **G** | .03 (.511) | .12 (.008)* | .03 (.535) | -.00 (.932) | -.02 (.616) | .10 (.021) | .08 (.058) | .07 (.103) | .05 (.218) | .05 (.239) |
| **CoS** | .04 (.400) | .12 (.007)* | .03 (.479) | -.02 (.717) | -.01 (.755) | .11 (.013) | .08 (.058) | .08 (.062) | .06 (.176) | .06 (.188) |
| **SI** | .01 (.780) | .12 (.008)* | .02 (.605) | -.03 (.554) | .00 (.933) | .10 (.019) | .10 (.027) | .08 (.061) | .07 (.117) | .06 (.162) |

*Note*: Partial correlations between general intelligence scores and condition-specific brain-average coupling values (measure-specific averages of coupling values from all brain regions) during eight different fMRI conditions controlled for age, gender, handedness, and in-scanner head motion. Significant associations passing the Bonferroni-corrected threshold (four comparisons) are marked with an asterisk (* = $p < .0125$). PL = path length; G = communicability; CoS = cosine similarity; SI = search information; RES = resting state; EMO = emotion processing task; GAM = gambling task; LAN = language task; MOT = motor task; REL = relational processing task; SOC = social cognition task; WM = working memory task; TG = task-general (average measure-specific whole-brain coupling across all seven task fMRI conditions); CG = condition-general (average measure-specific whole-brain coupling across all eight fMRI conditions).

**Supplementary Table S5**

Assessment of significant differences in prediction performance between the total of nine 5-fold cross-validated prediction models.

| | RES | EMO | GAM | LAN | MOT | REL | SOC | WM |
|---|---|---|---|---|---|---|---|---|
| **EMO** | RES > EMO<br>$\Delta|r|$ = .05<br>*p* = .288 | | | | | | | |
| **GAM** | GAM > RES<br>$\Delta|r|$ = .01<br>*p* = .487 | GAM > EMO<br>$\Delta|r|$ = .06<br>*p* = .270 | | | | | | |
| **LAN** | RES > LAN<br>$\Delta|r|$ = .03<br>*p* = .361 | LAN > EMO<br>$\Delta|r|$ = .03<br>*p* = .406 | GAM > LAN<br>$\Delta|r|$ = .04<br>*p* = .344 | | | | | |
| **MOT** | RES > MOT<br>$\Delta|r|$ = .11<br>*p* = .095 | EMO > MOT<br>$\Delta|r|$ = .05<br>*p* = .241 | GAM > MOT<br>$\Delta|r|$ = .12<br>*p* = .074 | LAN > MOT<br>$\Delta|r|$ = .08<br>*p* = .148 | | | | |
| **REL** | RES > REL<br>$\Delta|r|$ = .06<br>*p* = .271 | EMO > REL<br>$\Delta|r|$ = .00<br>*p* = .491 | GAM > REL<br>$\Delta|r|$ = .06<br>*p* = .253 | LAN > REL<br>$\Delta|r|$ = .03<br>*p* = .393 | REL > MOT<br>$\Delta|r|$ = .05<br>*p* = .238 | | | |
| **SOC** | SOC > RES<br>$\Delta|r|$ = .04<br>*p* = .355 | SOC > EMO<br>$\Delta|r|$ = .10<br>*p* = .183 | SOC > GAM<br>$\Delta|r|$ = .03<br>*p* = .364 | SOC > LAN<br>$\Delta|r|$ = .07<br>*p* = .235 | **SOC > MOT**<br>**$\Delta|r|$ = .15**<br>***p* = .047*** | SOC > REL<br>$\Delta|r|$ = .10<br>*p* = .167 | | |
| **WM** | WM > RES<br>$\Delta|r|$ = .08<br>*p* = .234 | WM > EMO<br>$\Delta|r|$ = .13<br>*p* = .106 | WM > GAM<br>$\Delta|r|$ = .07<br>*p* = .239 | WM > LAN<br>$\Delta|r|$ = .11<br>*p* = .139 | **WM > MOT**<br>**$\Delta|r|$ = .19**<br>***p* = .022*** | WM > REL<br>$\Delta|r|$ = .13<br>*p* = .092 | WM > SOC<br>$\Delta|r|$ = .03<br>*p* = .370 | |
| **ALL** | ALL > RES<br>$\Delta|r|$ = .09<br>*p* = .165 | ALL > EMO<br>$\Delta|r|$ = .15<br>*p* = .053 | ALL > GAM<br>$\Delta|r|$ = .08<br>*p* = .168 | ALL > LAN<br>$\Delta|r|$ = .12<br>*p* = .075 | **ALL > MOT**<br>**$\Delta|r|$ = .20**<br>***p* = .004*** | **ALL > REL**<br>**$\Delta|r|$ = .15**<br>***p* = .038*** | ALL > SOC<br>$\Delta|r|$ = .05<br>*p* = .316 | ALL > WM<br>$\Delta|r|$ = .01<br>*p* = .466 |

*Note*: Significant differences in prediction performance between all possible pairs of models (5-fold internal cross-validation) were assessed by comparing the absolute difference in prediction performance (correlation coefficients between predicted and observed intelligence scores) between two models trained on the observed intelligence scores ($\Delta|r_{observed}|$; reported in table) to the absolute difference in prediction performance when the same two models were trained on the permuted intelligence scores ($\Delta|r_{permuted}|$). *P*-values indicating significant model differences are marked with an asterisk (* = *p* < .05; $\Delta|r_{permuted}|$ was greater than $\Delta|r_{observed}|$ for less than 50/1.000 permutations). RES = resting state; EMO = emotion processing task; GAM = gambling task; LAN = language task; MOT = motor task; REL = relational processing task; SOC = social cognition task; WM = working memory task; ALL = all task FMRI conditions.

**Supplementary Table S6**

Differences in brain-average SC-FC coupling between fMRI conditions in the lockbox sample (HCP 232).

| FMRI condition 1 | FMRI condition 2 | *p*-value |
|---|---|---|
| RES | EMO | < .001* |
| RES | GAM | < .001* |
| RES | LAN | < .001* |
| RES | MOT | < .001* |
| RES | REL | < .001* |
| RES | SOC | < .001* |
| RES | WM | < .001* |
| EMO | GAM | < .001* |
| EMO | LAN | < .001* |
| EMO | MOT | < .001* |
| EMO | REL | < .001* |
| EMO | SOC | < .001* |
| EMO | WM | < .001* |
| GAM | LAN | < .001* |
| GAM | MOT | .991 |
| GAM | REL | .891 |
| GAM | SOC | < .001* |
| GAM | WM | < .001* |
| LAN | MOT | < .001* |
| LAN | REL | < .001* |
| LAN | SOC | < .001* |
| LAN | WM | < .001* |
| MOT | REL | .378 |
| MOT | SOC | < .001* |
| MOT | WM | < .001* |
| REL | SOC | < .001* |
| REL | WM | < .001* |
| SOC | WM | .068 |

*Note*: Brain-average SC-FC coupling strength significantly differed between fMRI conditions in the lockbox sample (ANOVA: F(7,1848) = [519.6], $p < .001$). Significant group differences are marked with an asterisk (Tukey's HSD procedure: * = $p < .05$; uncontrolled for multiple comparisons). RES = resting state; EMO = emotion processing task; GAM = gambling task; LAN = language task; MOT = motor task; REL = relational processing task; SOC = social cognition task; WM = working memory task.

**Supplementary Table S7**

Differences in brain-average SC-FC coupling between fMRI conditions in the replication sample (AOMIC PIOP1).

| FMRI condition 1 | FMRI condition 2 | *p*-value |
|---|---|---|
| RES | WM | < .001* |
| RES | EMO-A | < .001* |
| RES | EMO-M | < .001* |
| RES | FAC | < .001* |
| RES | GEN | < .001* |
| WM | EMO-A | < .001* |
| WM | EMO-M | < .001* |
| WM | FAC | < .001* |
| WM | GEN | < .001* |
| EMO-A | EMO-M | < .001* |
| EMO-A | FAC | < .001* |
| EMO-A | GEN | .221 |
| EMO-M | FAC | < .001* |
| EMO-M | GEN | < .001* |
| FAC | GEN | < .001* |

*Note*: Brain-average SC-FC coupling strength significantly differed between fMRI conditions in the AOMIC PIOP1 sample (ANOVA: $F(5,750) = [353.0]$, $p < .001$). Significant group differences are marked with an asterisk (Tukey's HSD procedure: * = $p < .05$; uncontrolled for multiple comparisons). RES = resting state; WM = working memory task; EMO-A = emotion anticipation task; EMO-M = emotion matching task; FAC = face perception task; GEN = gender stroop task.

**Supplementary Table S8**

Differences in brain-average SC-FC coupling between fMRI conditions in the replication sample (AOMIC PIOP2).

| FMRI condition 1 | FMRI condition 2 | *p*-value |
|---|---|---|
| RES | WM | < .001* |
| RES | EMO-M | < .001* |
| RES | STO | < .001* |
| WM | EMO-M | < .001* |
| WM | STO | < .001* |
| EMO-M | STO | < .001* |

*Note*: Brain-average SC-FC coupling strength significantly differed between fMRI conditions in the AOMIC PIOP2 sample (ANOVA: $F(3,716) = [209.8]$, $p < .001$). Significant group differences are marked with an asterisk (Tukey's HSD procedure: * = $p < .05$; uncontrolled for multiple comparisons). RES = resting state; WM = working memory task; EMO-M = emotion matching task; STO = stop signal task.

**Supplementary Table S9**

Differences in brain-average SC-FC coupling between coupling measures in the lockbox sample (HCP 232).

| **Measure 1** | **Measure 2** | ***p*-value** |
|---|---|---|
| PL | G | < .001* |
| PL | CoS | < .001* |
| PL | SI | < .001* |
| G | CoS | < .001* |
| G | SI | < .001* |
| CoS | SI | < .001* |

*Note*: Brain-average SC-FC coupling strength significantly differed between coupling measures in the lockbox sample (ANOVA: $F(3,924) = [542.0]$, $p < .001$). Significant group differences are marked with an asterisk (Tukey's HSD procedure: * = $p < .05$; uncontrolled for multiple comparisons). PL = path length; G = communicability; CoS = cosine similarity; SI = search information.

**Supplementary Table S10**

Differences in brain-average SC-FC coupling between coupling measures in the replication sample (AOMIC PIOP1).

| Measure 1 | Measure 2 | *p*-value |
|---|---|---|
| PL | G | < .001* |
| PL | CoS | < .001* |
| PL | SI | < .001* |
| G | CoS | .071 |
| G | SI | < .001* |
| CoS | SI | < .001* |

*Note*: Brain-average SC-FC coupling strength significantly differed between coupling measures in the AOMIC PIOP1 sample (ANOVA: $F(3,500) = [239.3]$, $p < .001$). Significant group differences are marked with an asterisk (Tukey's HSD procedure: * = $p < .05$; uncontrolled for multiple comparisons). PL = path length; G = communicability; CoS = cosine similarity; SI = search information.

**Supplementary Table S11**

Differences in brain-average SC-FC coupling between coupling measures in the replication sample (AOMIC PIOP2).

| **Measure 1** | **Measure 2** | ***p*-value** |
|---|---|---|
| PL | G | < .001* |
| PL | CoS | < .001* |
| PL | SI | < .001* |
| G | CoS | < .001* |
| G | SI | < .001* |
| CoS | SI | < .001* |

*Note*: Brain-average SC-FC coupling strength significantly differed between coupling measures in the AOMIC PIOP2 sample (ANOVA: $F(3,716) = [375.5]$, $p < .001$). Significant group differences are marked with an asterisk (Tukey's HSD procedure: * = $p < .05$; uncontrolled for multiple comparisons). PL = Path Length; G = Communicability; CoS = Cosine Similarity; SI = Search Information.

**Supplementary Table S12**

Relationship between general intelligence and brain-average SC-FC coupling during all fMRI conditions in the lockbox sample (HCP232).

| SC-FC coupling measure | FMRI condition *r* (*p*) | | | | | | | | | |
|---|---|---|---|---|---|---|---|---|---|---|
| | RES | EMO | GAM | LAN | MOT | REL | SOC | WM | TG | CG |
| **PL** | .08 (.239) | .06 (.360) | .04 (.512) | -.08 (.206) | -.00 (.972) | .01 (.855) | .01 (.901) | .13 (.054) | .02 (.738) | .03 (.613) |
| **G** | .04 (.526) | .05 (.498) | -.02 (.815) | -.14 (.035) | -.06 (.398) | -.01 (.897) | -.05 (.456) | .06 (.383) | -.04 (.551) | -.03 (.660) |
| **CoS** | .04 (.560) | .03 (.638) | .00 (.989) | -.16 (.019) | -.05 (.428) | -.00 (.965) | -.04 (.514) | .08 (.222) | -.04 (.583) | -.03 (.682) |
| **SI** | .02 (.765) | .01 (.905) | .08 (.257) | -.12 (.068) | -.01 (.836) | .02 (.787) | .01 (.862) | .13 (.059) | .01 (.866) | .01 (.848) |

*Note*: Partial correlations between general intelligence scores and condition-specific brain-average coupling values (measure-specific averages of coupling values from all brain regions) during eight different fMRI conditions controlled for age, gender, handedness, and in-scanner head motion in the lockbox sample. Significant associations passing the Bonferroni-corrected threshold (four comparisons) are marked with an asterisk (* = $p < .0125$). PL = path length; G = communicability; CoS = cosine similarity; SI = search information; RES = resting state; EMO = emotion processing task; GAM = gambling task; LAN = language task; MOT = motor task; REL = relational processing task; SOC = social cognition task; WM = working memory task; TG = task-general (average measure-specific whole-brain coupling across all seven task fMRI conditions); CG = condition-general (average measure-specific whole-brain coupling across all eight fMRI conditions).

**Supplementary Table S13**

Relationship between general intelligence and brain-average SC-FC coupling during all fMRI conditions in the replication sample (AOMIC PIOP1).

| SC-FC coupling measure | **FMRI condition** *r* (*p*) | | | | | | | |
|---|---|---|---|---|---|---|---|---|
| | RES | WM | EMO-A | EMO-M | FAC | GEN | TG | CG |
| **PL** | .13 (.153) | -.02 (.851) | -.14 (.118) | .04 (.668) | -.13 (.139) | -.01 (.919) | -.07 (.461) | -.02 (.801) |
| **G** | .13 (.160) | -.02 (.835) | -.16 (.082) | .10 (.297) | -.11 (.208) | -.01 (.922) | -.05 (.557) | -.02 (.863) |
| **CoS** | .15 (.091) | -.04 (.700) | -.17 (.054) | .09 (.333) | -.17 (.059) | -.00 (.975) | -.08 (.385) | -.03 (.781) |
| **SI** | .12 (.172) | -.00 (.948) | -.07 (.469) | .07 (.420) | -.13 (.166) | -.03 (.748) | -.03 (.755) | .00 (.974) |

*Note*: Partial correlations between general intelligence scores and condition-specific brain-average coupling values (measure-specific averages of coupling values from all brain regions) during six different fMRI conditions controlled for age, gender, handedness, and in-scanner head motion in the AOMIC PIOP1 sample. Significant associations passing the Bonferroni-corrected threshold (four comparisons) are marked with an asterisk (* = $p < .0125$). PL = path length; G = communicability; CoS = cosine similarity; SI = search information; RES = resting state; WM = working memory task; EMO-A = emotion anticipation task; EMO-M = emotion matching task; FAC = face perception task; GEN = gender stroop task; TG = task-general (average measure-specific whole-brain coupling across all five task fMRI conditions); CG = condition-general (average measure-specific whole-brain coupling across all six fMRI conditions).

**Supplementary Table S14**

Relationship between general intelligence and brain-average SC-FC coupling during all fMRI conditions in the replication sample (AOMIC PIOP2).

| **SC-FC coupling measure** | **FMRI condition** *r* (*p*) | | | | | |
|---|---|---|---|---|---|---|
| | RES | WM | EMO-M | STO | TG | CG |
| **PL** | -.02 (.799) | -.09 (.234) | -.01 (.941) | -.02 (.771) | -.045 (.541) | -.04 (.584) |
| **G** | -.02 (.802) | -.06 (.437) | .02 (.777) | -.00 (.954) | -.02 (.832) | -.02 (.826) |
| **CoS** | -.03 (.740) | -.04 (.580) | .02 (.840) | .00 (.969) | -.01 (.909) | -.01 (.848) |
| **SI** | -.01 (.883) | -.02 (.776) | .00 (.967) | .01 (.884) | -.00 (.977) | -.00 (.962) |

*Note*: Partial correlations between general intelligence scores and condition-specific brain-average coupling values (measure-specific averages of coupling values from all brain regions) during four different fMRI conditions controlled for age, gender, handedness, and in-scanner head motion in the AOMIC PIOP2 sample. Significant associations passing the Bonferroni-corrected threshold (four comparisons) are marked with an asterisk (* = $p < .0125$). PL = path length; G = communicability; CoS = cosine similarity; SI = search information; RES = resting state; WM = working memory task; EMO-M = emotion matching task; STO = stop signal task; TG = task-general (average measure-specific whole-brain coupling across all three task fMRI conditions); CG = condition-general (average measure-specific whole-brain coupling across all four fMRI conditions).

**Supplementary Table S15**

Performance when predicting intelligence from brain region-specific SC-FC coupling in the lockbox sample (HCP232).

| **FMRI condition** | **Prediction performance** | | |
|---|---|---|---|
| | *r* | $R^2$ | *p* |
| Resting state (RES) | .06 | < .01 | .232 |
| Emotion processing (EMO) | .02 | < .01 | .400 |
| Gambling (GAM) | .06 | < .01 | .145 |
| Language (LAN) | .25 | .06 | .006* |
| Motor (MOT) | .06 | < .01 | .214 |
| Relational processing (REL) | .06 | < .01 | .144 |
| Social cognition (SOC) | .06 | < .01 | .164 |
| Working memory (WM) | .17 | .03 | .018* |
| All tasks (ALL) | .21 | .04 | .003* |

*Note:* Intelligence predictions in the lockbox sample from brain region-specific coupling data during single tasks were based on the Basic NMA Model, while intelligence prediction from combined region-specific coupling data across all task fMRI conditions was realized with the Expanded NMA Model (see Methods). Significance of the prediction performance was assessed with non-parametric permutation tests. *P*-values indicating significant associations are marked with an asterisk (* = $p < .05$).

**Supplementary Table S16**

Performance when predicting intelligence from brain region-specific SC-FC coupling in the replication sample (AOMIC PIOP1).

| FMRI condition | Prediction performance | | |
|---|---|---|---|
| | *r* | $R^2$ | *p* |
| Resting state (RES) | .11 | .01 | .164 |
| Working memory (WM) | -.12 | - | .973 |
| Emotion anticipation (EMO-A) | -.01 | - | .515 |
| Emotion matching (EMO-M) | -.23 | - | .994 |
| Face perception (FAC) | -.32 | - | .998 |
| Gender stroop (GEN) | -.08 | - | .886 |
| All tasks (ALL) | -.11 | - | .880 |

*Note:* Intelligence predictions in the AOMIC PIOP1 sample from brain region-specific coupling data during single tasks were based on the Basic NMA Model, while intelligence prediction from combined region-specific coupling data across all task fMRI conditions was realized with the Expanded NMA Model (see Methods). Significance of the prediction performance was assessed with non-parametric permutation tests. *P*-values indicating significant associations are marked with an asterisk (* = $p < .05$).

**Supplementary Table S17**

Performance when predicting intelligence from brain region-specific SC-FC coupling in the replication sample (AOMIC PIOP2).

| FMRI condition | Prediction performance | | |
|---|---|---|---|
| | *r* | $R^2$ | *p* |
| Resting state (RES) | -.08 | - | .873 |
| Working memory (WM) | -.05 | - | .756 |
| Emotion matching (EMO-M) | -.04 | - | .726 |
| Stop signal (STO) | -.08 | - | .865 |
| All tasks (ALL) | -.04 | - | .717 |

*Note:* Intelligence predictions in the AOMIC PIOP2 sample from brain region-specific coupling data during single tasks were based on the Basic NMA Model, while intelligence prediction from combined region-specific coupling data across all task fMRI conditions was realized with the Expanded NMA Model (see Methods). Significance of the prediction performance was assessed with non-parametric permutation tests. *P*-values indicating significant associations are marked with an asterisk (* = $p < .05$).

**Supplementary Table S18**

Performance when predicting intelligence from brain region-specific SC-FC coupling in the cross-sample model generalization test in the lockbox sample (HCP532 → HCP232).

| FMRI condition | Prediction performance | | |
|---|---|---|---|
| | *r* | $R^2$ | *p* |
| Resting state (RES) | .15 | .02 | < .001* |
| Emotion processing (EMO) | .00 | < .001 | .422 |
| Gambling (GAM) | .05 | < .001 | .059 |
| Language (LAN) | .19 | .04 | < .001* |
| Motor (MOT) | .08 | .01 | < .001* |
| Relational processing (REL) | .13 | .02 | .002* |
| Social cognition (SOC) | .23 | .05 | < .001* |
| Working memory (WM) | .11 | .01 | .031* |
| All tasks (ALL) | .20 | .04 | < .001* |

*Note:* Intelligence predictions in the cross-sample model generalization test (HCP532 → HCP232) from brain region-specific coupling data during single tasks were based on the Basic NMA Model, while intelligence prediction from combined region-specific coupling data across all task fMRI conditions was realized with the Expanded NMA Model (see Methods). Significance of the prediction performance was assessed with non-parametric permutation tests. *P*-values indicating significant associations are marked with an asterisk (* = $p < .05$).

**Supplementary Table S19**

Assessment of significant differences in prediction performance between the total of nine models of the cross-sample model generalization test in the lockbox sample (HCP532 → HCP232).

| | RES | EMO | GAM | LAN | MOT | REL | SOC | WM |
|---|---|---|---|---|---|---|---|---|
| **EMO** | **RES > EMO**<br>**Δ\|r\| = .15**<br>**$p$ < .001*** | | | | | | | |
| **GAM** | **RES > GAM**<br>**Δ\|r\| = .11**<br>**$p$ = .015*** | GAM > EMO<br>Δ\|r\| = .04<br>$p$ = .122 | | | | | | |
| **LAN** | LAN > RES<br>Δ\|r\| = .03<br>$p$ = .341 | **LAN > EMO**<br>**Δ\|r\| = .18**<br>**$p$ = .001*** | **LAN > GAM**<br>**Δ\|r\| = .14**<br>**$p$ = .014*** | | | | | |
| **MOT** | RES > MOT<br>Δ\|r\| =. 07<br>$p$ = .075 | **MOT > EMO**<br>**Δ\|r\| = .08**<br>**$p$ = .005*** | MOT > GAM<br>Δ\|r\| = .04<br>$p$ = .158 | LAN > MOT<br>Δ\|r\| = .10<br>$p$ = .054 | | | | |
| **REL** | RES > REL<br>Δ\|r\| = 0.2<br>$p$ = .336 | **REL > EMO**<br>**Δ\|r\| = .13**<br>**$p$ = .006*** | REL > GAM<br>Δ\|r\| = .08<br>$p$ = .065 | LAN > REL<br>Δ\|r\| = .06<br>$p$ = .228 | REL > MOT<br>Δ\|r\| = .05<br>$p$ = .204 | | | |
| **SOC** | SOC > RES<br>Δ\|r\| = .07<br>$p$ = .177 | **SOC > EMO**<br>**Δ\|r\| = .22**<br>**$p$ < .001*** | **SOC > GAM**<br>**Δ\|r\| = .18**<br>**$p$ = .003*** | SOC > LAN<br>Δ\|r\| = .04<br>$p$ = .327 | **SOC > MOT**<br>**Δ\|r\| = .14**<br>**$p$ = .018*** | SOC > REL<br>Δ\|r\| = .10<br>$p$ = .108 | | |
| **WM** | RES > WM<br>Δ\|r\| = .05<br>$p$ = .236 | **WM > EMO**<br>**Δ\|r\| = .10**<br>**$p$ = .041*** | WM > GAM<br>Δ\|r\| = .06<br>$p$ = .165 | LAN > WM<br>Δ\|r\| = .08<br>$p$ = .163 | WM > MOT<br>Δ\|r\| = .02<br>$p$ = .362 | REL > WM<br>Δ\|r\| = .02<br>$p$ = .374 | SOC > WM<br>Δ\|r\| = .12<br>$p$ = .077 | |
| **ALL** | ALL > RES<br>Δ\|r\| = .05<br>$p$ = .248 | **ALL > EMO**<br>**Δ\|r\| = .20**<br>**$p$ < .001*** | **ALL > GAM**<br>**Δ\|r\| = .15**<br>**$p$ = .001*** | ALL > LAN<br>Δ\|r\| = .01<br>$p$ = .444 | **ALL > MOT**<br>**Δ\|r\| = .12**<br>**$p$ = .013*** | All > REL<br>Δ\|r\| = .07<br>$p$ = .148 | SOC > ALL<br>Δ\|r\| = .03<br>$p$ = .354 | ALL > WM<br>Δ\|r\| = .09<br>$p$ = .100 |

*Note*: Significant differences in prediction performance between all possible pairs of models (cross-sample model generalization test; HCP532 → HCP232) were assessed by comparing the absolute difference in prediction performance (correlation coefficients between predicted and observed intelligence scores) between two models trained on the observed intelligence scores (Δ|$r_{observed}$|; reported in table) to the absolute difference in prediction performance when the same two models were trained on the permuted intelligence scores (Δ|$r_{permuted}$|). *P*-values indicating significant model differences are marked with an asterisk (* = $p$ < .05; Δ|$r_{permuted}$| was greater than Δ|$r_{observed}$| for less than 50/1.000 permutations). RES = resting state; EMO = emotion processing task; GAM = gambling task; LAN =

language task; MOT = motor task; REL = relational processing task; SOC = social cognition task; WM = working memory task; ALL = all task fMRI conditions.

**Supplementary Table S20**

Performance when predicting intelligence from brain region-specific SC-FC coupling in the cross-sample model generalization test in the replication sample (HCP532 → AOMIC PIOP1).

| **FMRI condition** | **Prediction performance** | | |
|---|---|---|---|
| | *r* | $R^2$ | *p* |
| Resting state (RES) | .04 | < .01 | .216 |
| Working memory (WM) | .15 | .02 | < .001* |
| Emotion matching (EMO-M) | .12 | .01 | < .001* |

*Note:* Intelligence predictions in the cross-sample model generalization test (HCP532 → AOMIC PIOP1) from brain region-specific coupling data during single tasks were based on the Basic NMA Model, while intelligence prediction from combined region-specific coupling data across all task fMRI conditions was realized with the Expanded NMA Model (see Methods). Significance of the prediction performance was assessed with non-parametric permutation tests. *P*-values indicating significant associations are marked with an asterisk (* = $p < .05$). Note that the cross-sample model generalization test of the Expanded NMA Model was not applicable due to differences in tasks applied during fMRI assessment.

**Supplementary Table S21**

Performance when predicting intelligence from brain region-specific SC-FC coupling in the cross-sample model generalization test in the replication sample (HCP532→ AOMIC PIOP2).

| FMRI condition | Prediction performance | | |
|---|---|---|---|
| | *r* | $R^2$ | *p* |
| Resting state (RES) | .05 | < .01 | .006* |
| Working memory (WM) | -.00 | - | .545 |
| Emotion matching (EMO-M) | .04 | - | .007* |

*Note:* Intelligence predictions in the cross-sample model generalization test (HCP532 → AOMIC PIOP2) from brain region-specific coupling data during single tasks were based on the Basic NMA Model, while intelligence prediction from combined region-specific coupling data across all task fMRI conditions was realized with the Expanded NMA Model (see Methods). Significance of the prediction performance was assessed with non-parametric permutation tests. *P*-values indicating significant associations are marked with an asterisk (* = $p < .05$). Note that the cross-sample model generalization test of the Expanded NMA Model was not applicable due to differences in tasks applied during fMRI assessment.

**Supplementary Table S22**

Differences in brain-average SC-FC coupling between fMRI conditions (control analysis; based on frame length 176).

| FMRI condition 1 | FMRI condition 2 | *p*-value |
|---|---|---|
| RES | EMO | < .001* |
| RES | GAM | < .001* |
| RES | LAN | < .001* |
| RES | MOT | < .001* |
| RES | REL | < .001* |
| RES | SOC | < .001* |
| RES | WM | < .001* |
| EMO | GAM | < .001* |
| EMO | LAN | < .001* |
| EMO | MOT | < .001* |
| EMO | REL | < .001* |
| EMO | SOC | < .001* |
| EMO | WM | < .001* |
| GAM | LAN | < .001* |
| GAM | MOT | .859 |
| GAM | REL | .648 |
| GAM | SOC | < .001* |
| GAM | WM | 1.000 |
| LAN | MOT | < .001* |
| LAN | REL | < .001* |
| LAN | SOC | < .001* |
| LAN | WM | < .001* |
| MOT | REL | 1.00 |
| MOT | SOC | < .001* |
| MOT | WM | .972 |
| REL | SOC | < .001* |
| REL | WM | .868 |
| SOC | WM | < .001* |

*Note*: Brain-average SC-FC coupling strength based on shortened frame lengths (176) significantly differed between fMRI conditions (ANOVA: F(7,4248) = [542,07], $p < .001$). Significant group differences are marked with an asterisk (Tukey's HSD procedure: * = $p < .05$; uncontrolled for multiple comparisons). RES = resting state; EMO = emotion processing task; GAM = gambling task; LAN = language task; MOT = motor task; REL = relational processing task; SOC = social cognition task; WM = working memory task.

**Supplementary Table S23**

Performance when predicting intelligence from brain region-specific SC-FC coupling based on shortened frame lengths (176).

| FMRI condition | Prediction performance | | |
|---|---|---|---|
| | *r* | $R^2$ | *p* |
| Resting state (RES) | .19 | .04 | .001* |
| Emotion processing (EMO) | n/a | n/a | n/a |
| Gambling (GAM) | .20 | .04 | .001* |
| Language (LAN) | .15 | .02 | .002* |
| Motor (MOT) | .06 | < .01 | .108 |
| Relational processing (REL) | .12 | .01 | .014* |
| Social cognition (SOC) | .22 | .05 | .001* |
| Working memory (WM) | .17 | .03 | .004* |
| All tasks (ALL) | .25 | .06 | < .001* |

*Note:* Control analysis: all scan lengths were shortened to a frame length of 176 (matching the shortest task: emotion processing task) prior to computing the region-specific SC-FC coupling values. The Basic NMA Model was used to predict intelligence based on brain region-specific coupling obtained during single tasks, while the Expanded NMA Model was used to predict intelligence from task-combined region-specific coupling (see Methods). *P*-values indicating significant associations are marked with an asterisk (* = $p < .05$, non-parametric permutation tests).

**Supplementary Table S24**

FMRI conditions in the replication samples (AOMIC PIOP1 & AOMIC PIOP2)[5].

| | Condition | Description | Scan present | | Frames per run | | Run duration (min) | |
|---|---|---|---|---|---|---|---|---|
| | | | PIOP1 | PIOP2 | PIOP1 | PIOP2 | PIOP1 | PIOP2 |
| **1** | Resting state (RES) | Participants' eyes are open and focused on a fixation cross. | ✓ | ✓ | 480 | 240 | 6:00 | 8:00 |
| **2** | Working memory (WM) | Visual working memory task asking the participant whether the orientation of a bar had changed or not. | ✓ | ✓ | 162 | 162 | 5:24 | 5:24 |
| **3** | Emotion anticipation (EMO-A) | Presentation of a cue (reliable in 80% of cases) followed by a negatively valenced or neutral image measuring processes related to emotional anticipation and curiosity. | ✓ | x | 200 | x | 6:40 | x |
| **4** | Emotion matching (EMO-M) | Matching of emotional faces (anger/fear). | ✓ | ✓ | 135 | 135 | 4:30 | 4:30 |
| **5** | Face perception (FAC) | Passive watching of short video clips presenting dynamic facial expressions (anger, contempt, joy, pride, or a neutral condition). | ✓ | x | 330 | x | 4:07.5 | x |
| **6** | Gender stroop (GEN) | Face-gender variant of the Stroop task where different male or female faces paired with either a corresponding or opposite label are presented. This task measures processes related to cognitive control or conflict. | ✓ | x | 245 | x | 8:10 | x |
| **7** | Stop signal (STO) | Response inhibition task asking participants to quickly decide whether a male or female face is presented except when an auditory stop signal is heard. | x | ✓ | x | Scan length differs between subjects: 210-250 frames; median frames 224 | x | 7:00 – 8:20<br>Median 7:28 |

*Note:* Data from resting state and a total of six task fMRI conditions were acquired for the AOMIC PIOP1 and AOMIC PIOP2 samples. Besides scan length and application of multi-slice acceleration, the image acquisition parameters did not differ between fMRI conditions.

**Detailed Description of Similarity Measures and Communication Measures**

**Similarity Measures**

Similarity measures describe the conformity of regional structural connectivity profiles and are computed based on the SC matrix. They are ultimately represented in similarity matrices where each entry illustrates how the structural connections of brain region $i$ resemble the structural connections of brain region $j$. Note that the respective regional structural connectivity profiles are defined by matrix columns. For the computation of the similarity measure (cosine similarity; CoS) from each individual SC matrix, no further information about plausible communication strategies is included. Therefore, this operationalization serves as a 'baseline' measure of SC-FC coupling. Measure descriptions are adapted from Popp et al.[6] and Zamani Esfahlani et al.[7].

**Cosine Similarity**

Cosine similarity measures the resemblance between two brain regions' connectivity profiles (matrix columns) based on their orientation in an $N - 1$ dimensional connectivity space, where $N$ is the number of brain regions, i.e., 358. We calculated the cosine similarity of the angle between two vectors $x = [x_1, \dots, x_N]$ and $y = [y_1, \dots, y_N]$ as $CoS_{xy} = \frac{x \cdot y}{\|x\| \cdot \|y\|}$, where vectors are region-specific connectivity profiles for every possible pair of brain regions[8].

**Communication Measures**

Communication measures reflect the how easy it is for two brain regions to communicate based on the underlying structural connections and a proposed signaling conceptualization (i.e., communication model like shortest path routing, navigation, random walks)[9,10]. This 'ease of communication' is subsequently represented in communication matrices. Particularly, each individual weighted SC matrix was transformed into three communication matrices representing three distinct communication measures.

**Path Length**

Path length is a metric of how easily signals can be transmitted between two regions via their shortest path, as longer paths are more susceptible to noise, have longer delays in transmission and are energetically more costly[9,11]. In a network, each edge is associated with a cost $C$ (difficulty of traversing) and for weighted networks, this cost can be assessed by transforming the weight $\omega$

of each edge into a measure of length through $C = \omega^{-1}$. The shortest path between a pair of nodes (source node $s$ and target node $t$) is the sequence of edges $\pi_{s\rightarrow t} = \{A_{si}, A_{ij}, \ldots, A_{mt}\}$ minimizing the sum $C_{si} + C_{ij} + \cdots + C_{mt}$ (where $C_{si}$ is the cost of traversing the edge between region $s$ and $i$) and $i, j$ and $m$ are nodes along the shortest path.

**Communicability**

Communicability assumes that neural signaling unfolds as a diffusive broadcasting process, supposing that information can flow along all possible walks between two brain regions[12,13]. It can be characterized as the weighted sum of all walks of all lengths between two respective regions[14], where an edge is the connection between two brain regions and a walk is a sequence of traversed edges. This measure accounts for all possible connections between regions but includes walk lengths ($l_w$) and penalizes the contribution of walks with increased lengths. For weighted networks, SC matrices ($A$) are first normalized as $A' = D^{-1/2}AD^{-1/2}$, where $D$ is the degree diagonal matrix[15]. The normalized matrix is then exponentiated to calculate the communicability as $G = e^{A'}$ or $G = \sum_{wl=0}^{\infty} \frac{A'^{l_w}}{l_w!}$, where each walk is inversely proportional to its length thus 1-step walks contribute $\frac{A'^1}{1!}$, 2-step walks $\frac{A'^2}{2!}$ and so on.

**Search Information**

Search information is a measure of network navigability without global knowledge[16,17]. It is related to the probability that a random walker will travel between two nodes via their shortest path (i.e., the path connecting two nodes via fewest intermediate stations/nodes). This probability increases with an expanding number of paths that are available for a certain communication process to take place[9,16]. Given the shortest path between brain regions $s$ (source node) and $t$ (target node): $\pi_{s\rightarrow t} = \{s, i, j, \ldots, l, m, t\}$, the probability of accessing this shortest path is expressed as $F(\pi_{s\rightarrow t}) = f_{si} \times f_{ij} \times \ldots \times f_{lm} \times f_{mt}$, where $f_{ij} = \frac{A_{ij}}{\sum_j A_{ij}}$ and $i, j, l$ and $m$ are nodes along the shortest path. The information that is then required to access the shortest path from $s$ to $t$ is $SI(\pi_{s\rightarrow t}) = \log_2 [F(\pi_{s\rightarrow t})]$[16].

As search information and path length depict difficulty of communication as opposed to ease of communication, respective communication matrices were transformed to also reflect ease of communication and therefore allow easier interpretation of analysis results (for details refer to Popp et al.[6]). Further, communication matrices for search information are asymmetric, suggesting that the ease of communication between region $i$ to $j$ doesn't necessarily equal the ease of communication between region $j$ and $i$[18]. To ensure comparability between similarity and communication measures, respective individual communication matrices were symmetrized (for summary of similarity and communication measures see Table 2).

**Detailed Description of Replication Samples (AOMIC PIOP1 & AOMIC PIOP2)**

To further assess the generalizability of our study findings, all analyses were also conducted in two completely independent datasets from the Amsterdam Open MRI Collection (AOMIC)[5] deviating in e.g., data acquisition and preprocessing as well as in the general intelligence measure. Specifically, we used data from the AOMIC PIOP1 sample and the AOMIC PIOP2 sample. Data for the AOMIC PIOP1 sample was acquired on the 'Achieva' version of a Philips 3T scanner, while data for the PIOP2 sample was obtained from the 'Achieva sStream Version' of the Philips 3T scanner while using a 32-channel head coil. For both datasets, one diffusion-weighted scan was included in the analysis pipeline (median TR = 7387 ms; TE = 86 ms; 2-mm isotropic voxel-resolution; b = 1000 s/mm$^2$; 32 directions/shell) and the resulting structural imaging data were preprocessed similarly to the HCP data except white matter fibers were modeled using a maximum spherical harmonics of order six. Regarding fMRI (BOLD) scans, the AOMIC PIOP1 included a resting-state and five task fMRI conditions (Supplementary Table S24). The AOMIC PIOP2 sample entails fMRI scans during resting state and three task fMRI conditions (Supplementary Table S24). For the AOMIC PIOP1 sample, fMRI data from the resting-state scan and one additional task (face perception task) were acquired with multi-slice acceleration (TR = 750 ms; TE = 28 ms; 3-mm isotropic voxel resolution; flip angle = 60°; multiband acceleration factor 3), while all other fMRI scans in in the AOMIC PIOP1 sample and the AOMIC PIOP2 sample applied sequential acquisition (TR = 2000 ms; TE = 28 ms; 3-mm isotropic voxel resolution; flip angle = 76.1°). FMRI data were downloaded in the minimally preprocessed form, and we applied the same preprocessing steps as used in the main sample. Additional details about the imaging data acquisition for the two replication samples can be found in Snoek et al.[5]. To approximate general intelligence, the sum score of the 36 item version (set II) of the Raven's Advanced Progressive Matrices Test[19,20] was used. After excluding subjects based on the same criteria applied in the main sample (missing demographic, behavioral or neuroimaging data; excessive in-scanner head motion), 126 participants remained in the AOMIC PIOP1 sample (70 female; 112 right-handed; mean age =

22.24 years; age range = 18.25-26 years) and 180 participants remained in the AOMIC PIOP2 sample (103 female; 160 right-handed; mean age = 21.91 years; age range = 18.25-25.5 years).
